# A Murine Model of *Mycobacterium abscessus* Encapsulated in Alginate-Beads : Advancing Toward a Chronic Infection Model

**DOI:** 10.64898/2026.04.24.720549

**Authors:** Maya Rima, Aurélie Chauffour, Régis Tournebize, Corentin Poignon, Sarah Wong, Thi Cuc Mai, Maria Bitar, Romina Mehrabdollahi, Noel Zahr, Thibaud Coradin, Alexandra Aubry, Nicolas Veziris

## Abstract

**Background:** The increasing incidence of *Mycobacterium abscessus* (*M. abscessus*) lung infections, together with its intrinsic multidrug resistance, highlights the need for new therapeutic regimens. However, the lack of a reliable chronic infection model in immunocompetent mice limits preclinical evaluation.

**Methods:** To mimic the bronchial environment of infected patients, we evaluated the effect of encapsulating *M. abscessus* in alginate beads on infection progression in BALB/cJRJ mice following intratracheal inoculation, compared with intranasal infection using non-encapsulated bacteria. The impact of dexamethasone treatment (DEX) was also assessed. Bacterial loads in lungs, spleen, liver, and kidneys were quantified over time in untreated and antibiotic-treated mice. Lung inflammation was evaluated by measuring IFN-γ and TNF-α levels. *In vitro*, the activity of imipenem and bedaquiline was assessed against free or alginate-encapsulated *M. abscessus*.

**Results:** Compared with intranasal infection, intratracheal infection with alginate-encapsulated bacteria resulted in slower pulmonary clearance and greater extrapulmonary dissemination. DEX further enhanced these features, reducing lung clearance, increasing dissemination, and amplifying lung inflammation. Bedaquiline showed no effect, whereas imipenem efficacy depended on treatment timing. For both drugs, alginate encapsulation reduced *in vitro* antibacterial activity.

**Conclusion:** This model represents a step toward a chronic *M. abscessus* infection model characterized by moderate lungs clearance, extrapulmonary dissemination, and pronounced inflammatory responses. Reduced antibiotic activity against alginate-encapsulated bacteria may more accurately predict treatment efficacy in humans than activity measured against free bacteria.

## INTRODUCTION

Nontuberculous mycobacteria (NTM) are responsible for a wide spectrum of opportunistic infections in humans, with steadily increasing global incidence [1]. Among them, *Mycobacterium abscessus* (*M. abscessus*) is one of the most frequently isolated from patients with pulmonary infections. The clinical presentation is that of a chronic pulmonary disease, accompanied by systemic inflammation in the most severe cases. Due to its intrinsic and acquired multidrug resistance, it represents a substantial health issue for patients at risk, those with underlying lung diseases or compromised immune systems [2, 3]. Given the limitations of the current treatments (prolonged duration, limited efficacy and adverse effects), there is an urgent need for novel anti-*M. abscessus* drugs.

Preclinical murine studies are crucial for the development of novel drug candidates and have enabled the evaluation of numerous compounds against mycobacteria [4-6]. Among *M. abscessus* preclinical models described in the literature, some reproduce acute infection, limiting the efficacy testing to the early active replicative phase [7-9]. Chronic infection models with sustained infection are mainly established in genetically modified immunocompromised mice (nude, SCID…), which are not only costly but also do not accurately reproduce the pathology observed in humans [7, 10] Following a series of workshops, a consensus paper defined the key characteristics of a chronic murine model of *M. abscessus* infection: immunocompetent mice, increasing or sustained pulmonary bacterial load for at least 28 days, and features mimicking chronic human pulmonary disease [11].

The critical limitation of the murine models arises from the rapid clearance of *M. abscessus* in immunocompetent mice, hindering the establishment of a chronic infection [12, 13]. To address this issue, Riva *et al*., explored strategies to enhance bacterial evasion of host defenses, specifically by encapsulating *M. abscessus* in agar beads, limiting physical clearance and promoting bacterial retention within the lung airways [14]. However, both in this first study and another study by Malcolm *et al*., bacterial load stability beyond the two first weeks after infection and extrapulmonary dissemination was not confirmed in all experiments [14, 15].

On the other hand, corticosteroid-induced immunosuppression in immunocompetent mice has emerged as a promising approach, enabling bacterial proliferation in the lungs while maintaining sufficient host response to drive tissue pathology [13, 16, 17]. Although this approach resulted in chronic lung infection with an increasing bacterial load, the use of genetically modified C3HeB/FeJ mice in most studies, with limited supplies, is a limitation of this model.

Moreover, the antibiotic activity demonstrated in some murine models contrasts with the very limited activity of antibiotic regimens in humans [7, 17]. Thus, there is a need for a model predicting more accurately antibiotic efficacy.

In this study, our objectives were to (i) set up a model of chronic *M. abscessus* infection characterized by a continuous increase in bacterial load in lungs, or at least stability over one month, using an affordable immunocompetent mouse model, and (ii) identify a control antibiotic effective against *M. abscessus* in this model. To achieve these goals, we evaluated the effects of *M. abscessus* encapsulation in alginate beads and corticosteroids treatment, specifically dexamethasone (DEX), on the development of chronic infection in mice. We also tested the effects of imipenem (IMP) and bedaquiline (BDQ) in this model.

## METHODS

### Ethics statement

The study was approved by ethics committee n°5 Charles Darwin, Pitié-Salpêtrière Hospital (Paris, France) and authorized by the French Ministry of Higher Education and Research under the number APAFIS #31933-2021040214468842 v11. The animal facility was accredited (license number D75-13-08). Personnel were trained in accordance with French and Europeans recommendations. Experimental designs followed the guidelines ARRIVE 2.0 [18].

### Antibiotics and corticosteroid

IMP (Imipinem/Cilastatine, arrow®, 500 mg/500 mg, laboratoire arrow) doses were prepared daily in saline solution (0.9% NaCl, Versol). BDQ (bedaquiline fumarate, 17221182, Fisher) was prepared in 20% of acidified (2-Hydroxypropyl)-β-cyclodextrine (HPCD) (H107, Sigma-Aldrich) and conserved at 4°C for the duration of the experiment. DEX was purchased from Sigma-Aldrich (D1756). Daily treatment doses of DEX were pre-weighed, stored at -20°C, and freshly prepared in phosphate-buffered saline (1X PBS, Sigma-Aldrich).

### Mycobacterial strain and growth media

We used the rough variant of *M. abscessus* ATCC 19977, kindly provided by Jean-Louis Hermann (Infection and Inflammation, INSERM U1173, UVSQ). The broth medium used was 7H9 (217310, BD) supplemented with 10% OADC (211886, BD), and 0.05% of Tween 80 (2002-C, Euromedex). Bacterial enumeration was performed on 7H11 (212203, BD) supplemented with 0.4% charcoal (C9157, Sigma-Aldrich), and antibiotics (Mycobacteria Selectatab, MS24, MastGroup). MICs of IMP and BDQ determined by microdilution were 16 and 0.125 mg/L, respectively.

### Preparation of *M. abscessus* suspension encapsulated in alginate beads

Bacteria were encapsulated in alginate beads (∼100 µm/bead) using the Nisco encapsulation system (VARJ30, Nisco Engineering AG, Zurich, Switzerland) (S1). The encapsulated suspension was stored at 4°C for no longer than one week.

### Time-kill kinetics of BDQ and IMP on *M. abscessus* encapsulated or non-encapsulated in alginate beads

Bacterial suspensions of encapsulated and non-encapsulated bacteria (∼ 10^5^ CFU/mL) were prepared in 7H9 medium supplemented with 10% OADC and 0.05% Tween 80 and exposed to antibiotics. BDQ and IMP were tested at 0.5x, 1x, 2x, 4x, and 8x their respective MICs. To compensate the IMP instability, an equivalent amount of IMP was added every 24h [19]. Cultures were incubated at 30°C under agitation (120 rpm). Bacteria were quantified by plating on 7H11 (30°C, 5 days) after 48, 72, and 96h of incubation.

### Murine experiments

#### Experiment 1: comparison of intranasal infection with M. abscessus suspension and intratracheal infection with M. abscessus encapsulated in alginate beads (S2)

BALB/cJRJ mice (female, 4-5 weeks old, Janvier breeding center, Le Genest-Saint-Isle, France) were intratracheally inoculated with encapsulated bacteria (50 µL). Treatments with IMP and BDQ started at D1 and continued for 7, 14, or 30 days. For comparative purposes, 20 mice were infected intranasally with non-encapsulated bacteria. Lung bacterial loads and dissemination to spleen, liver, and kidneys were assessed. Lung inflammation was assessed by measuring IFN-γ and TNF-α levels.

#### Experiment 2: evaluation of corticosteroid treatment impact on evolution of intratracheal infection with M. abscessus encapsulated in alginate beads (S3)

DEX was administered once-daily from one week before infection to mice infected intranasally or intratracheally with non-or encapsulated bacteria, respectively. IMP treatment started at D7 post-infection for one or four weeks. Bacterial loads in lungs, liver, spleen and kidneys, and lung cytokines (IFN-γ and TNF-α) were quantified.

### Graphs and statistical analyses

Analyses and graphs were done using GraphPad Prism 10.5.0 for windows (GraphPad Software, USA). All groups were compared to D1, treated groups to their corresponding untreated controls, and CFU in untreated group were compared across time points. We used one-way ANOVA, with Kruskal-Wallis test for multiple comparisons. Differences were considered significant when *p*-value <0.05.

## RESULTS

### Effect of M. abscessus encapsulation on the establishment of chronic infection in mice

#### CFU counts

Mice intranasally infected with non-encapsulated bacteria showed a significant decrease of lung bacterial loads as early as the second week of infection and half of the mice had a bacillary load beyond the limit of detection after 1 month (Figure 1A). No extrapulmonary dissemination was detected.

**Figure 1.**
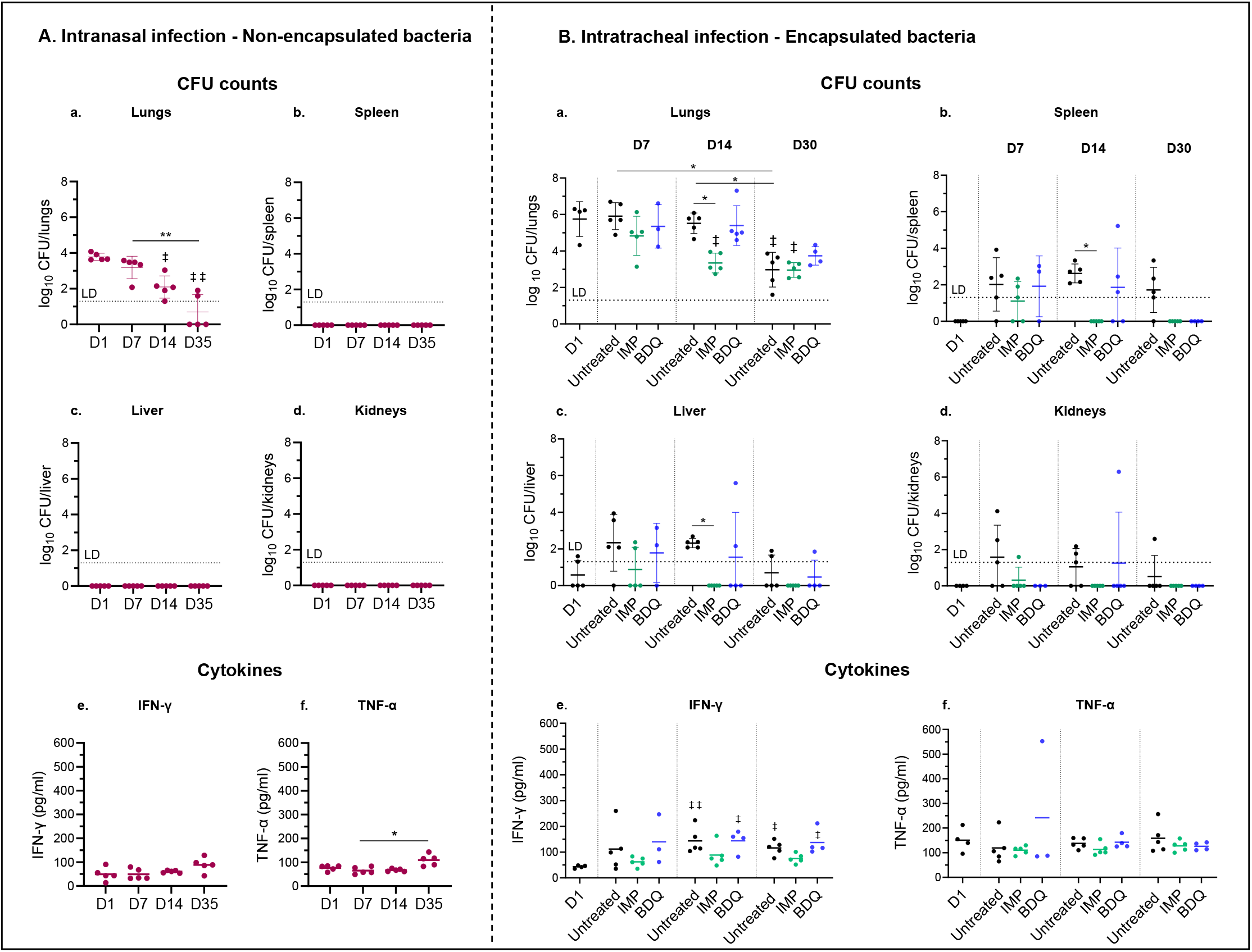
*M. abscessus* burden in organs of mice intranasally infected with non-encapsulated bacteria (4.4 log10 CFU/mouse) (A) or intratracheally infected with encapsulated bacteria (4.3 log_10_ CFU/mouse) (B) untreated or treated daily (5/7d) with IMP (100 mg/kg, BID, subcutaneous) or BDQ (25 mg/kg/d, orally) for 1 (D7), 2 (D14) or 4 weeks (D30) (LD: limit of detection). Cytokines (IFN-γ and TNF-α) quantified in the lung homogenates. Results are presented as individual values, with the group mean ± SD. Statistically significant differences were determined by one-way ANOVA with Kruskal-Wallis test for multiple comparisons between D1 and all other groups (‡, *p*-value < 0.05; ‡‡, *p*-value < 0.01), and both between the treated groups and the corresponding untreated one and within the untreated groups over time (*, *p*-value < 0.05; **, *p*-value < 0.01). n=5 mice/group unless reduced by unexpected mortality.

Differently, mice intratracheally infected with encapsulated bacteria, had stable lung CFU counts at D7 and D14, before showing a significant decrease at D30 (3.0 ± 1.0 log_10_ CFU/lungs vs 5.8 ± 1.0 log_10_ CFU/lungs at D1) (Figure 1B). Extrapulmonary dissemination appeared from D1 in the liver (0.7 ± 0.9 log_10_ CFU/liver) and D7 in the spleen (2.0 ± 1.5 log_10_ CFU/spleen), remaining stable thereafter. The kidneys showed the lowest bacterial burden, together with marked variability.

#### Inflammatory response

In mice intranasally infected, TNF-α increased only at D35 while IFN-γ levels remained stable over time (Figure 1A). In mice intratracheally infected, TNF-α levels were significantly higher than the intranasal group, particularly at D14 (*p*-value = 0.015), but without significant variation over time (Figure 1B). In contrast, IFN-γ levels significantly increased during infection.

### Effect of corticosteroid treatment on the establishment of chronic infection in mice

#### CFU counts

In intranasally infected mice, DEX treatment showed no effect compared with the first experiment. Similar bacterial clearance was observed with also 3/5 mice showing negative lung cultures at D35 (Figure 2A). A limited extrapulmonary dissemination was observed with 0 to 3 mice culture-positive at different time-points.

**Figure 2.**
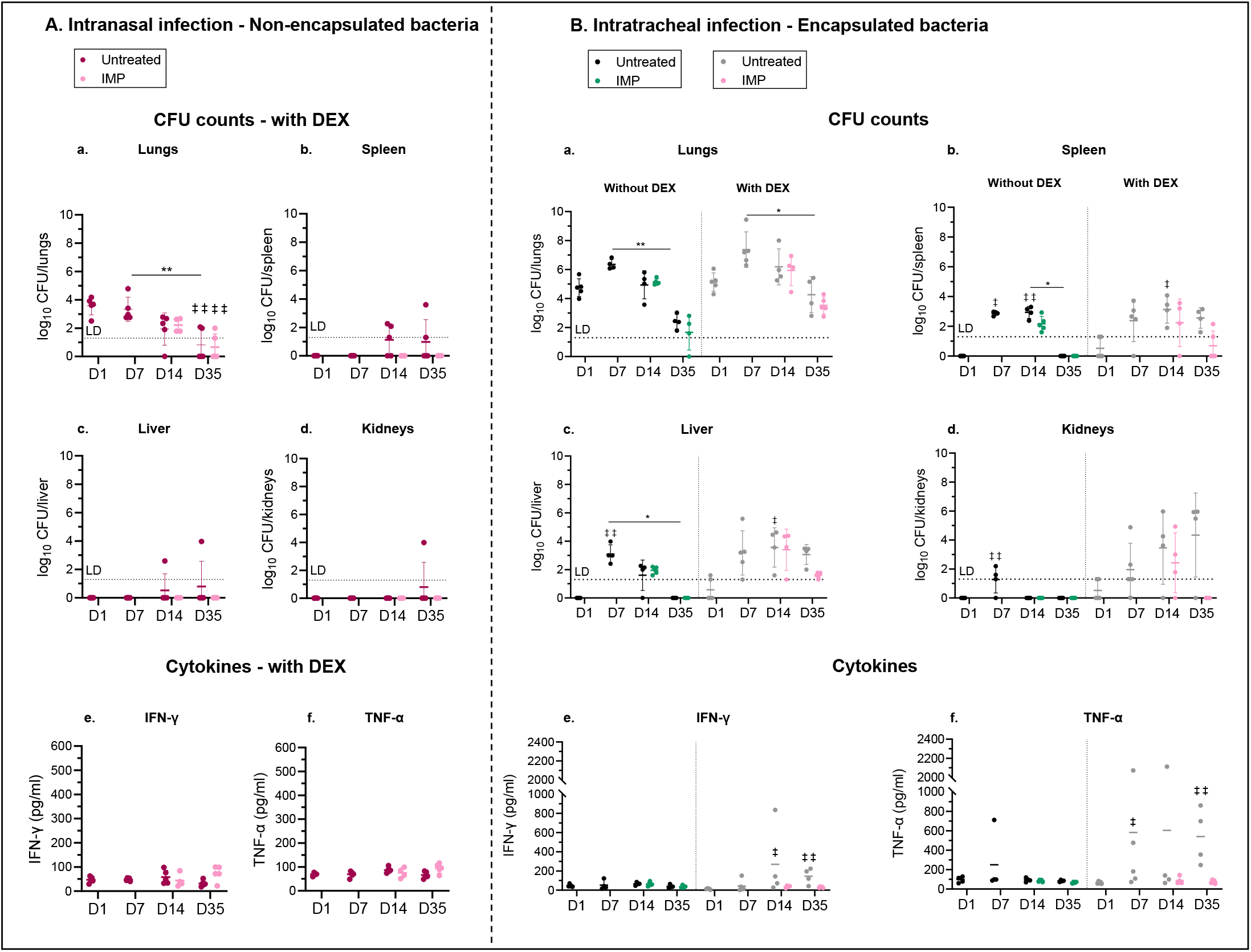
*M. abscessus* burden in organs of mice intranasally infected with non-encapsulated bacteria without DEX treatment (4.4 log_10_ CFU/mouse) (a) or intratracheally infected with encapsulated bacteria (4.6 log_10_ CFU/mouse) (b), treated or not with DEX (5 mg/kg/d, subcutaneous, 5/7d), and treated or not with IMP (100 mg/kg, BID, subcutaneous, 5/7d, starting at D7) (LD: limit of detection). Cytokines (IFN-γ and TNF-α) quantified in the lung homogenates. Results are presented as individual values, with the group mean ± SD. Statistically significant differences were determined by one-way ANOVA with Kruskal-Wallis test for multiple comparisons between D1 and all other groups (‡, *p*-value < 0.05; ‡‡, *p*-value < 0.01), and both between the treated groups and the corresponding untreated one and within the untreated groups over time (*, *p*-value < 0.05; **, *p*-value < 0.01). n=5 mice/group unless reduced by unexpected mortality.

In intratracheally infected groups, the CFU kinetics in non DEX-treated mice confirmed those observed in the first experiment (Figure 2B). DEX showed significantly slowed bacterial clearance with only 1 log_10_ CFU/lungs of reduction at D35 compared to D1, whereas the non-treated group achieved nearly 2 log_10_ CFU/lungs of reduction. Bacteria became detectable from D7 in spleen (2.4 ± 1.4 log_10_ CFU/spleen) and liver (3.2 ± 1.6 CFU log_10_ CFU/liver) in DEX-treated mice. In this latter group, bacterial load remained stable overtime in both organs, whereas it declined by D35 in non-DEX-treated mice. Interestingly, an important bacterial dissemination in kidneys was observed in DEX-treated mice: bacteria became detectable from D7 and gradually increased, reaching 4.3 ± 2.9 log_10_ CFU/kidneys at D35, comparable to lung levels. As in first experiment, large variations in CFU were observed.

#### Inflammatory response

In mice infected intranasally, DEX had no significant effect except at D35 (Figure 2A). In mice infected intratracheally, DEX significantly affected cytokines profiles (Figure 2B). While cytokines amount remained stable over time in the non-treated group, an increase was observed in the DEX-treated one, becoming significant from D7 and D14 for TNF-α and IFN-γ, respectively. These levels were sustained until D35, with TNF-α level significantly higher (*p*-value = 0.012) than that in DEX-treated, intranasally infected mice.

### Identification of a control antibiotic for the M. abscessus murine model

When given from D1 in non-DEX-treated mice, IMP reduced lung bacterial loads, particularly at D7 and D14, with decreases of almost 1 and 2 log_10_ CFU/lungs compared to untreated groups (Figure 1B). It also prevented extrapulmonary dissemination, with no detectable cultivable bacteria from D14 and thereafter. A similar reduction of inflammatory biomarkers was observed, particularly for IFN-γ, showing a noticeable yet non-significant reduction relative to the untreated group, especially at D14. In contrast, BDQ treatment had no significant effect on bacterial burden or inflammatory biomarkers, except in the spleen at D30. The high CFU variability in the kidneys limits the interpretation of IMP and BDQ treatment outcomes in this organ.

To assess the impact of treatment timing, IMP was administered one-week post-infection in Exp.2, instead of 1-day post-infection, while BDQ was excluded due to its ineffectiveness in Exp.1. Similar to mice infected intranasally with non-encapsulated bacteria, IMP had no effect on bacterial burden or inflammatory cytokines production in the lungs of either DEX or non-DEX-treated mice intratracheally infected with encapsulated bacteria (Figure 2A,B). However, IMP reduced extrapulmonary dissemination, particularly in DEX-treated mice, with spleen and liver bacterial loads decreased by 2 log_10_ CFU/spleen and 1.5 log_10_ CFU/liver at D35 compared to the untreated groups (Figure 2B). IMP was also effective in the kidneys of DEX-treated mice, with no detectable bacteria at D35. Interestingly, a similar trend was observed in lung inflammatory biomarkers of DEX-treated mice, showing a noticeable yet non-significant reduction vs untreated groups.

### Time-kill kinetics of BDQ and IMP on *M. abscessus* encapsulated or non-encapsulated in alginate beads

At the highest concentrations (4x and 8x MIC), BDQ was bacteriostatic against both encapsulated and non-encapsulated bacteria (Figure 3, a,c). However, CFU counts were consistently higher for encapsulated bacteria. This difference reached statistical significance at 2×MIC, where bacterial growth at 96h relative to the initial inoculum was significant for encapsulated bacteria (*p*-value = 0.007) but not for non-encapsulated bacteria (Table 1).

**Table 1.** Time-kill kinetics of BDQ against *M. abscessus* non-encapsulated or encapsulated in alginate beads. Results are expressed as means ± SD from three independent experiments. Starting inoculum: 5.8 ± 0.4 log_10_ CFU/mL.

| BDQ concentrations (µg/mL) |  | Comparison versus the starting inoculum<br>log<sub>10</sub> CFU/mL at 96h – log<sub>10</sub> CFU/mL at 0h |  |
| --- | --- | --- | --- |
|  |  | Non-encapsulated bacteria | Encapsulated bacteria |
| 0.5 x MIC | 0.0625 | +2.3 (± 0.3) | +2.2 (± 0.8) |
| MIC | 0.125 | +1.6 (± 0.3) | +2.0 (± 0.9) |
| 2 x MIC | 0.25 | +0.9 (± 0.7) | +1.7 (± 0.2) |
| 4 x MIC | 0.5 | +0.2 (± 0.2) | +0.9 (± 0.79) |
| 8 x MIC | 1 | -0.1 (± 0.3) | +0.4 (± 0.8) |

**Figure 3.**
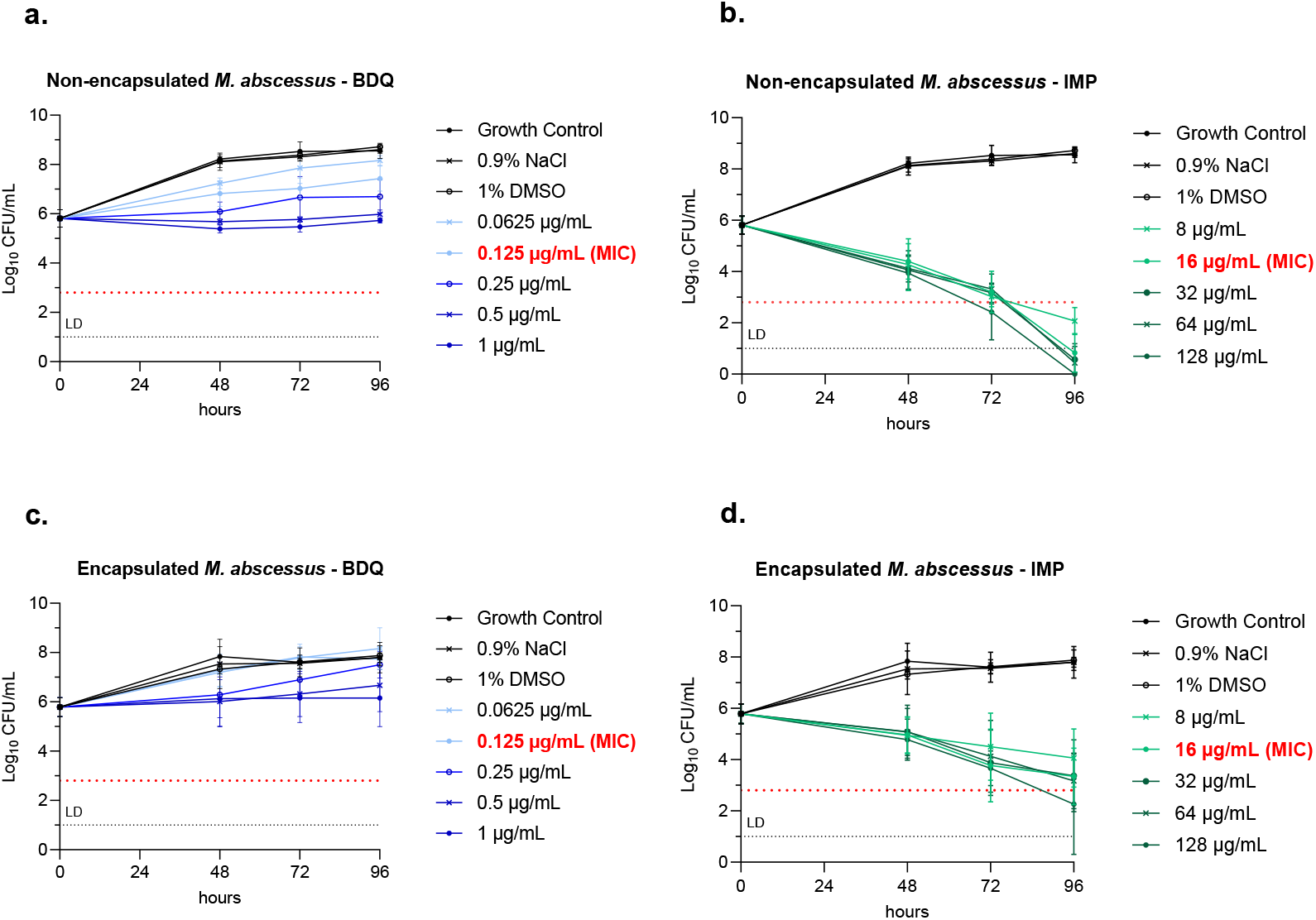
Time-kill kinetics of bedaquiline (BDQ) and imipenem (IMP) against *M. abscessus* non-encapsulated (a,b) or encapsulated (c,d) in alginate beads. MIC of BDQ: 0.125 µg/mL. MIC of IMP: 16 µg/mL. Red dashed line: bactericidal limit corresponding to 3 log_10_ CFU/mL of reduction relative to the inoculum. LD: limit of detection. Results are expressed as means ± SD from three independent experiments.

Regarding IMP, a pronounced dose-dependent bactericidal effect was observed in non-encapsulated bacteria at all tested concentrations, with a significant reduction in bacterial counts (> 3 log_10_ reduction) at 96h compared to the inoculum (Figure 3b). In encapsulated bacteria, a 3 log_10_ reduction was observed only at 128 µg/mL but did not reach statistical significance compared to the initial inoculum (Figure 3d). Overall, IMP showed reduced activity against encapsulated *M. abscessus*, with nearly >2 log_10_ difference compared to non-encapsulated bacteria (Table 2).

**Table 2.** Time-kill kinetics of IMP against *M. abscessus* non-encapsulated or encapsulated in alginate beads. Results are expressed as means ± SD from three independent experiments. Starting inoculum: 5.8 ± 0.4 log_10_ CFU/mL.

| IMP concentrations (µg/mL) |  | Comparison versus the starting inoculum<br>log<sub>10</sub> CFU/mL at 96h – log<sub>10</sub> CFU/mL at 0h |  |
| --- | --- | --- | --- |
|  |  | Non-encapsulated bacteria | Encapsulated bacteria |
| 0.5 x MIC | 8 | -3.8 (± 0.8) | -1.7 (± 0.8) |
| MIC | 16 | -5.0 (± 0.6) | -2.5 (± 0.7) |
| 2 x MIC | 32 | -5.3 (± 0.6) | -2.4 (± 1.1) |
| 4 x MIC | 64 | -5.4 (± 1.0) | -2.6 (± 0.9) |
| 8 x MIC | 128 | -5.9 ± 0.3 | -3.5 ± 1.6 |

## DISCUSSION

Encapsulating bacteria, as a potential barrier against host clearance, was initially developed to establish chronic lung infection with *Pseudomonas aeruginosa* (*P. aeruginosa*) [20]. When translated to *M. abscessus* using agar beads, this model achieved sustained lung colonization for up to 90 days with minimal spleen dissemination [14]. Here, we investigated the effect of *M. abscessus* encapsulation in alginate beads on infection progression and dissemination. Alginate was chosen due to its presence in the infected pulmonary environment, where it constitutes a major component of *P. aeruginosa* biofilm matrix in cystic fibrosis (CF) patients [21]. Moreover, *M. abscessus* is known to co-infect with *P. aeruginosa*, with nearly 80% of CF patients infected with *M. abscessus* also harboring *P. aeruginosa* [22]. Our objective was to mimic this infectious biofilm, which can alter the effect of antibiotics as demonstrated for aminoglycosides [23, 24]. Interestingly, while the intranasal infection with non-encapsulated bacteria resulted in a reduction of >1.5 log_10_ CFU/lungs at D14, pulmonary bacterial load in mice intratracheally infected with encapsulated bacteria remained stable for 14 days post-infection before showing a significant decrease at D30 (Figures 1A-B). Unlike Riva *et al*., these findings align with Malcolm *et al*., who infected C57Bl/6J mice with *M. abscessus* encapsulated in agar beads [15]. However, even in Riva *et al*., the CFU stability observed with *M. abscessus* was not reproduced with *M. massiliense* and *M. bolletii* [14]. Although encapsulating bacteria in alginate beads did not fully prevent clearance, it displayed promising features, indicating progress toward a persistent infection with extrapulmonary dissemination and pulmonary inflammation.

We then evaluated the combined effect of encapsulated bacteria and DEX-treatment on infection progression and dissemination. Beyond establishing a chronic model, steroids appear relevant since they represent a significant risk factor for NTM pulmonary infection, reported in one-third of patients on oral steroids and about half on inhaled steroids [25, 26].

Corticosteroid, particularly DEX, has been reported to promote chronic *M. abscessus* infection in mice exposed to aerosolized, non-encapsulated bacteria [13, 17]. However, our results showed no impact of DEX on infection progression in mice intranasally infected with non-encapsulated bacteria (Figure 2A), in contrast with Nicklas *et al*., who observed a continuous increase in lung bacterial loads for up to 4 weeks [17]. This difference may be explained by several factors. First, while C3HeB/FeJ mice are frequently utilized, we used BALB/cJRJ, which may account for the observed difference [27-29]. However, Maggioncalda *et al*., and Sriram *et al*., showed that this model could also be applied to BALB/c with 1-2 weeks of DEX treatment prior to infection [13, 16]. This leads us to reconsider other possible causes of this difference, such as the frequency of DEX-treatment. While DEX was administered 7 days/week in most studies, we treated mice 5 days/week. Finally, to verify the actual dose administered, we measured the concentration of the DEX solution (S4). Although the target was 0.5 mg/mL (5 mg/kg), it was found to be 0.3 mg/mL, 40% lower than intended. Since previous studies have mentioned that the DEX solution, as ours, was cloudy, we assume that DEX is not fully soluble in PBS. To ensure proper solubility in PBS, an hydrosoluble DEX reference might have been more appropriate than the one reported in the literature [13, 16, 28, 29] and will be used in our future experiments.

In mice infected with alginate-encapsulated bacteria, DEX-treatment impaired bacterial clearance in lungs and other organs, particularly in kidneys, where a high bacterial load was detected (Figure 2B). To the best of our knowledge, the combined effect of DEX and alginate beads-based infection has not been previously reported in the literature.

A significant increase in IFN-γ and TNF-α levels, indicative of pronounced pulmonary inflammation, was recorded predominantly in DEX-treated mice intratracheally infected with encapsulated bacteria. This response persisted until D35, with TNF-α levels remaining significantly higher than those in the intranasal groups. This aligns with clinical data showing systemic inflammation in patients with pulmonary *M. abscessus* or *M. avium*-complex infection, which is associated with increased mortality [30, 31].

Since the murine chronic *M. abscessus* model aims to evaluate novel drugs, incorporating an effective reference antibiotic for comparison purposes is primordial. In our beads-model, we tested IMP and BDQ, two antibiotics with promising activity in prior *M. abscessus* models [7, 29, 32]. Results showed no effect of BDQ (Figure 1B), in contrast to the literature using non-encapsulated *M. abscessus* [7, 29]. This difference may be attributed to encapsulation in alginate beads, which could interact with BDQ and potentially limit its effect by reducing bacterial accessibility, as seen with tobramycin against *P. aeruginosa* [33]. To investigate this hypothesis, we performed an *in vitro* time-to-kill assay (Figure 3). Consistent with the literature and with our *in vivo* results, BDQ exhibited a bacteriostatic effect [34, 35]. However, its activity appeared weaker against encapsulated bacteria. These findings may partially explain the lack of BDQ efficacy observed in our model. Since limited activity has also been reported in humans [36], this model may be more predictive of BDQ activity.

Regarding IMP, initiating treatment one-week post-infection reduced its effects in lungs and other organs of non-DEX-treated mice, showing that treatment efficacy in our beads-model depends on the timing of treatment initiation. In DEX-treated mice, IMP was ineffective in the lungs but reduced extrapulmonary dissemination and pulmonary inflammation (Figure 2B). This contrasts with other studies reporting high IMP efficacy in lungs when administered one week-post infection in DEX-treated models using non-encapsulated bacteria [17, 28]. Similar to BDQ, IMP showed reduced *in vitro* activity against encapsulated bacteria (∼ 2 log_10_ difference compared to non-encapsulated bacteria) (Figure 3). Overall, encapsulation in alginate beads reduced antibiotic activity and could explain why, despite their proven activity, imipenem-based regimens have a low success rate in humans [37].

These findings highlight the challenges of defining a standardized, effective reference antibiotic as a comparator in assessing the efficacy of novel candidates. They further emphasize that antibiotic responses vary with intrinsic model features.

In conclusion, we demonstrate that the *M. abscessus* beads murine model can be refined by incorporating alginate and, thus, more accurately mimic human lung pathology. Importantly, encapsulation in alginate beads reduced antibiotic activity, suggesting that this model may better predict antibiotic activity in humans. The introduction of steroid treatment was crucial, resulting in moderate decline in CFU, important extrapulmonary dissemination, and significant increase in inflammatory cytokine levels. Since this inflammation is clinically relevant and no cytokine increase was observed in the IMP-treated group, cytokines quantification, together with CFU counts, both hallmarks of human disease, may serve as valuable measurable biomarker for assessing treatment efficacy in this model.

### Financial support

This project has received funding from the Innovative Medicines Initiative 2 Joint Undertaking (JU) under grant agreement No [853932]. The JU receives support from the European Union’s Horizon 2020 research and innovation program and EFPIA.

